# Bacteria in honeybee crops are decoupled from those in floral nectar and bee mouths

**DOI:** 10.1101/2024.03.01.583024

**Authors:** Magdalena L. Warren, Kaoru Tsuji, Leslie E. Decker, Manabu Kishi, Jihoon Yang, Adina C. Howe, Tadashi Fukami

**Author notes:** These authors contributed equally to this work (Magdalena L. Warren, Kaoru Tsuji).

## Abstract

Bacteria in the honeybee gut are a well-recognized factor affecting bee health. However, the primary focus of this research has been the hindgut, while the crop, or honey stomach, is assumed to be dominated by environmentally acquired transient taxa that matter little to the bees. To evaluate this assumption, we examined bacterial taxa in the crop and mouth of *Apis mellifera* and *A. cerana japonica* foragers and in the nectar of *Prunus mume* flowers visited by the bees in the Minabe-Tanabe region of Japan. We found that in bacterial composition, the crop was distinct from both the mouth and the nectar, whereas mouth and nectar samples were indistinguishable. Furthermore, the crop remained similar in bacterial composition and diversity, while the mouth showed a sharp drop in alpha diversity and a large increase in beta diversity, from summer to winter. These results refute the conventional assumption, suggesting instead that the crop contains a conserved bacterial community largely distinct from environmental taxa. We also found that strains of a crop-associated species, *Apilactobacillus kunkeei*, could be season- and host species-specific. Together, these findings suggest that crop-associated bacterial communities should be studied further to better understand the relationship between honeybees and their gut bacteria.

## Introduction

Microbes in the honeybee gut are receiving increasing attention not only as a factor affecting the health of the agriculturally vital insects, but also as a model system for gaining basic understanding of gut-associated microbiota [1, 2]. One component of the honeybee gut is used for the transport of liquids. This organ, which is an expandable portion of the esophagus hereby termed the crop, but also known as the honey stomach or honey sac [3], contains species such as *Apilactobacillus kunkeei* and *Bombella apis* that may affect the survival and pathogen resistance of the foraging adults that host them and the larvae that the adults feed [1, 4–6]. However, microbes in the crop are not as well studied as those in the hindgut, which are generally considered more important to honeybee health [7, 8].

Given its function for temporary storage and transport of floral nectar [9], it seems reasonable to assume that the crop is occupied mainly by the transient microbes acquired from the environment via foraging of nectar [10, 11]. One way to investigate this assumption of environment–crop matching is to compare microbes in the crop of foraging bees and those in their mouth as well as in the floral nectar they forage for. If the assumption is true, the crop should be similar to both the mouth and the floral nectar in bacterial composition [4, 12]. To our knowledge, no study has directly compared bacteria in crop, mouth, and nectar for this purpose, leaving untested the assumption that the crop microbiota is dominated by environmentally derived transients.

Here we test the hypothesis that crop-associated bacteria show species composition that mirror those observed in mouth- and nectar-associated bacteria. To this end, we compare crop-, mouth-, and nectar-associated bacteria in samples that were collected simultaneously at the same locations. Our crop and mouth samples come from actively foraging adults of the introduced *Apis mellifera* and the native *A. cerana japonica* (hereafter *A. cerana*) collected in the Minabe-Tanabe region of Wakayama Prefecture in Japan. In this region, farmers use both species of honeybees to pollinate the winter-blooming Japanese apricot, *Prunus mume* [13]. Our nectar samples come from *P. mume* flowers collected in the winter near the hives of the *A. mellifera* and *A. cerana* bees that we caught for crop and mouth sampling.

## Materials and Methods

### Study sites

Japanese apricot orchards in the Minabe-Tanabe region are embedded within a mountainous countryside landscape that has recently become internationally recognized. In 2015, the Food and Agricultural Organization designated the region as a Globally Important Agricultural Heritage System for the agricultural practice in the region that embraces both human needs and biodiversity conservation. Many of the apricot orchards are located adjacent to natural and managed forests, which are thought to provide stable habitats for the native *A. cerana*. The two *Apis* species are the main pollinators of the apricot, which blooms from mid or late February to early or mid-March, during which few other flower-visiting insects are available [13]. In the winter, *A. mellifera* colonies are supplemented with a sugar solution, while *A. cerana* colonies are not. Although there may have been several different blooming species during our collections in winter and summer, our interest in nectar focused on the economically important *P. mume.* All our bee and nectar collection sites were located within this landscape, but at least 1.0 km away from their respective nearest neighboring collection site in each season (Fig. 1a, Table S1).

**Fig. 1.**
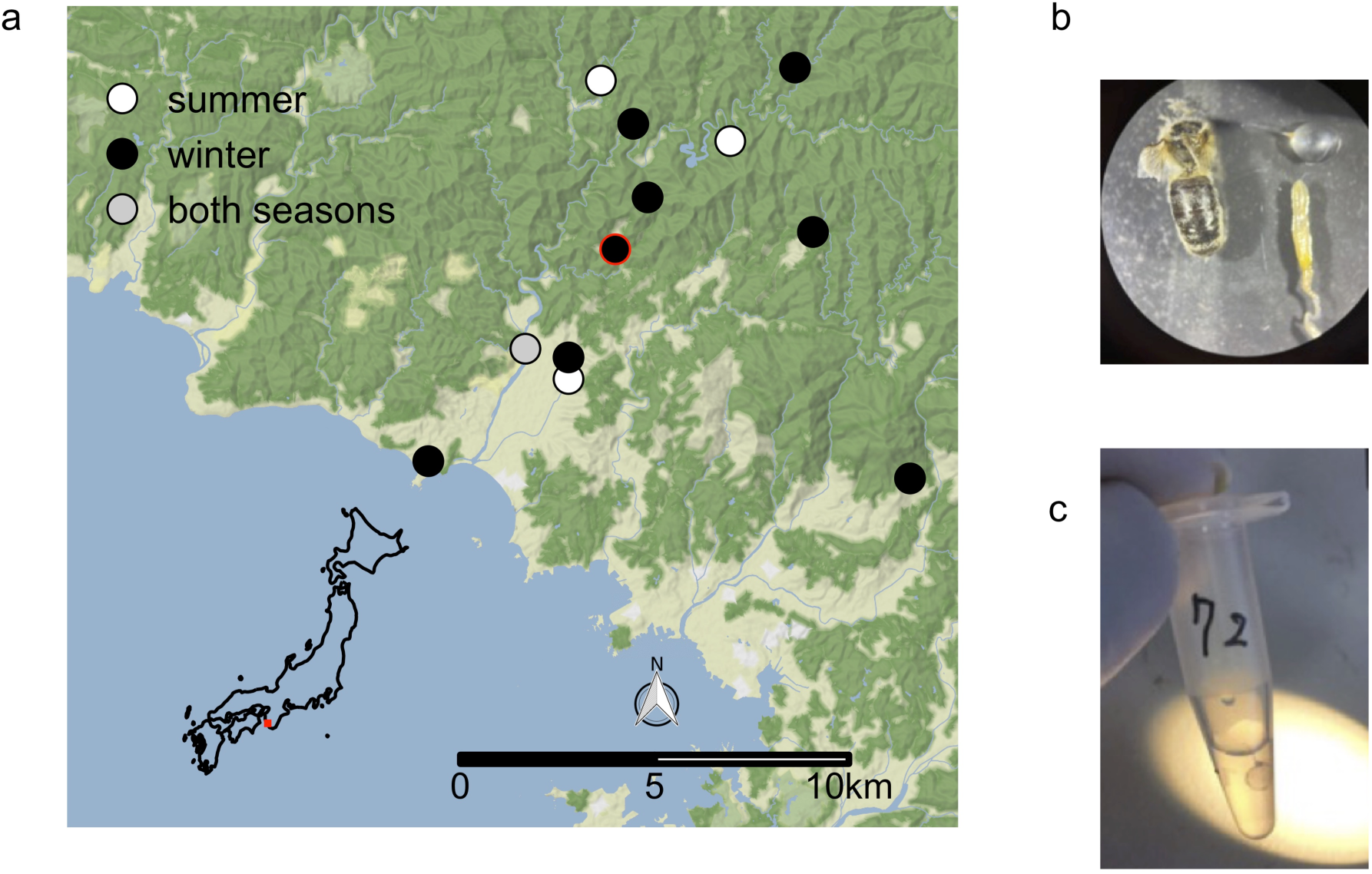
Collection sites and crop dissection. (**a**) Foraging adults of *Apis cerana* and *A. mellifera* were collected from a total of 12 sites (see Table S1) in or near Japanese apricot orchards in the summer of 2018 and the winter of 2019 in the Minabe-Tanabe region, the location of which is indicated by the red square on the map of Japan. Nectar samples were also collected from apricot flowers at each of the winter collection sites. Sites are labeled with circles colored by collection season (summer = white, winter = black, both seasons = grey). The location of the Japanese Apricot Laboratory is indicated by the red outline of the Higashi-Honjo site. (**b**) The crop was distinctly separated from the rest of the alimentary tract. (**c**) The crop of each honeybee was sterilely dissected and placed in an aliquot of sterile water

### Bee collection

A total of 12 *P. mume* orchards in the study region were visited in the summer of 2018 and the winter of 2019. Seventy-two and 149 honeybees were collected in the summer of 2018 and the winter of 2019, respectively, from hives in or near these apricot orchards (Table S1). At each collection site, there were about four to 10 *A. cerana* hives or 10 to 40 *A. mellifera* hives. Honeybees were collected as they flew near their hives located within or near the orchards. At orchards where the beekeepers maintained *A. ceran*a hives, *A. cerana* bees were collected near their hives whose locations varied from tens of meters to about a hundred meters from the orchard and were often situated at the edge of the local forest. The collected bees were individually placed in a sterile, sealed plastic vial immediately after the collection. The vials were kept for up to two hours in a cool box with ice packs in it for transport to the Japanese Apricot Laboratory located within 100 m from one of the collection sites (Higashi-Honjo) (Fig. 1a).

After we brought the collected bees into the laboratory in the Japanese Apricot Laboratory facility, the bees were kept cool in the box for up to two additional hours so that the bees would stay sluggish for ease of handling. The captured bees were then taken out of the vials, and their wings were fixated with adhesive tape to facilitate handling. The taped bees were made to drink approximately half of 20 µL of sterile 20% sucrose solution placed on the lid of a sterile 1.5-mL microcentrifuge tube. After the feeding, the remaining solution was placed in the microcentrifuge tube with the lid now closed. These solutions were later used to study the bacteria associated with the mouth. We refer to these solutions as mouth samples. As we had the bees extend their proboscis into the sterile sucrose solution, our method is likely to have captured mostly the bacteria from the surface of the proboscis rather than those present within the mandibles.

Immediately after the sampling of mouth-associated bacteria, we dissected the bees by gently pulling the abdomen using tweezers to expose the crop, making sure the crop did not touch other parts of the bee body. We then removed the crop from the bee body using another set of tweezers (Fig. 1b). Both tweezers were sterilized before each use by dipping in 70% ethanol and then flaming. Each crop sample was separately placed in 40 µL of sterile water within sterile microcentrifuge tubes (Fig. 1c) and homogenized using a sterile plastic pestle. All mouth and crop samples were immediately stored at −80°C until further processing.

We collected crop and mouth samples not just in winter during the *P. mume* flowering season, but also in summer, to determine whether crop and mouth differed in the way bacterial composition changed from summer to winter. Hives of *A. cerana*, many of which were made of carved tree trunks, were maintained by local beekeepers throughout the year in the region. In contrast, most *A. mellifera* hives in the region were transported elsewhere in the country while *P. mume* was not in bloom. However, some were kept locally throughout the year, allowing us to collect bees in both seasons. One site (Nishi-Honjo) was available for bee collection in both summer and winter, allowing bacterial taxa from the same site and hives to be compared between seasons. In addition, at two of our sites (Higashi-Honjo and Kamihaya), we were able to collect both *A. cerana* and *A. mellifera*, enabling comparison of the two species collected at the same locations (Table S1).

### Nectar collection

In addition to the mouth and crop samples, we also collected samples of *P. mume* floral nectar on the same days the bees were collected in the winter of 2019. At each of the nine orchards where we sampled bees that winter (Table S1), we collected 16 nectar samples (except at one site, Kamihaya, where we instead collected 24 nectar samples), for a total of 152 nectar samples. Each sample was collected by probing 10 flowers on a randomly selected branch of an individual *P. mume* tree with a sterile 0.5μl microcapillary tube and dispensing the collected nectar into a PCR tube containing 40 μL of PCR-quality sterile deionized water. The collected nectar samples were immediately placed in a box with ice packs during the transport to the laboratory. Within six hours, the nectar samples were brought back to the Japanese Apricot Laboratory. The samples were kept at −80°C until further processing for DNA extraction, except during a one-day transport to the laboratory at Kyoto University in Shiga, Japan, and another one-day transport from there to Stanford University in California, USA, during which the samples were kept at about −5 °C.

### Microbial DNA analysis

Using the nectar and bee samples obtained as above, we conducted microbial DNA extraction, bacterial amplicon sequencing, quantitative real-time PCR (qPCR) of 16S rRNA gene, and processing of the amplicon sequencing data (for details, see Text S1).

### Statistical analysis

All analyses were conducted using R version 4.0.4 [14]. The ASV_97_ dataset was analyzed with the phyloseq and vegan packages [15, 16]. Venn diagrams of sample types and seasons at the Genus level were created using the MicrobiotaProcess package [17, 18]. Differential abundances of bacteria agglomerated to the Genus level were explored with the DESeq2 package version 1.30.1 [19]. The taxa identified as significantly differentially abundant in the different contrasts (i.e. winter mouth vs. winter crop, winter nectar vs. winter crop, winter mouth vs. winter nectar, and summer crop vs. summer mouth; Fig S1) were aggregated. Of these taxa, those that were present in at least 50% of the sample type groups (Table S2) were further analyzed with a heatmap. The heatmap was created with the ComplexHeatmap package [20]. Differential abundance taxa phylogeny was made with a GTR model using the phangorn package [21]. Sample cladogram was based on a weighted UniFrac distance [22]. Alpha diversity (Shannon index) in each sample was calculated and compared by fitting a linear mixed effects model of Shannon index as the response and site as a random effect with 2000 parametric bootstraps [23]. Shannon-Weaver evenness was calculated using the evenness function from the microbiome package [24]. Permutation multivariate analysis of variance using a weighted UniFrac distance matrix [22] was used to test the effects of season, sample type, site, and host species on bacterial ASV_97_ composition. The beta diversity of relative abundance and absolute abundance samples was visualized with a Principal Coordinate Analysis, and individual sample distances from the centroid and the homogeneity of dispersion of the samples were calculated. We also did Multinomial Species Classification Method (CLAM) tests, using the clamtest package [25], to identify ASVs_97_ found more frequently in either winter or summer, those found frequently in both seasons, and those too rare to categorize as summer- or winter-associated, or generalists.

### Apilactobacillus kunkeei strain-level phylogenetic analysis

Fifty µL of each thawed honeybee crop sample were aliquoted onto De Man, Rogosa and Sharpe (MRS) agar (Sigma-Aldrich), spread thoroughly using a sterile loop, and incubated aerobically at 37°C for approximately 48 hours. Half of each single white colony was picked and placed in 10 µL of MilliQ (EMD Millipore, Burlington, MA, USA) water, and the other half was streaked onto a new MRS agar plate with a sterile loop. The 16S rRNA gene of these colonies was amplified using the universal primers 27F and 1492R in a 25 µL PCR amplification that included 12.5 µL of MyTaq Red Mix (Bioline), 8.75 µL of PCR-grade water, 1.25 µL each of the forward and reverse primers, 0.25 µL of dimethyl sulfoxide (DMSO), and 1 µL of each diluted colony. The resulting product was sequenced with Sanger sequencing, and colonies identified as *Apilactobacillus kunkeei* were once again isolated and placed in 10 µL of MilliQ (EMD Millipore) water. The variable segments of three housekeeping genes, *lepA*, *recG*, and *rpoB* were amplified using previously described primers [4, 26]. The 25 µL PCR amplification was made up of 11 µL of PCR-grade water, 12 µL of MyTaq Red Mix (Bioline), 0.5 µL of each forward and reverse primer, and 1 µL of the diluted colony. The annealing temperature for the PCR amplification was set to 47°C for *lepA* and *recG* and 60°C for *rpoB* for 30 seconds [4, 26]. The amplified PCR product was sequenced with Sanger sequencing. As previously described [27], the sequences were aligned with MAFFT [28] and trimmed with TrimAI [29] with a penalty for more than 50% gaps. Individual gene trees and the partitioned analysis for the multi-gene alignment were inferred using IQ-TREE [27, 30]. Statistical support for the phylogenies was calculated using 1000 replicates of both a Shimodaira-Hasegawa-like approximation likelihood test and ultrafast bootstrapping along with a Bayesian-like transformation of aLRT [27, 31, 32]. The partition model was selected by the IQ-TREE ModelFinder (Table S3) [27, 33]. *Apilactobacillus apinorum* Fhon13 (Table S4) [27] served as the outgroup for this phylogenetic tree. Outlier long branches were trimmed with TreeShrink [34].

## Results

### Crop–mouth–nectar comparisons

Among winter samples, we found more than seven times as many bacterial ASVs_97_ in the crop (a total of 655) as in the mouth and the nectar (a total of 77 and 81, respectively) (Fig. 2). Mean ASV_97_ diversity per sample was about six times higher in the crop (1.78) than in the mouth (0.37) and the nectar (0.36) (crop*–*mouth: Welch’s t = 17.1, *p-value* < 0.001; crop*–*nectar: Welch’s t = 15.8, *p-value* < 0.001). Similarly, mean evenness was about two times higher in the crop (0.74) than in the mouth (0.42) and the nectar (0.38) (crop*–*mouth: Welch’s t = 6.8, *p-value* < 0.001; crop*–*nectar: Welch’s t = 7, *p-value* < 0.001) (Fig. 3).

**Fig. 2.**
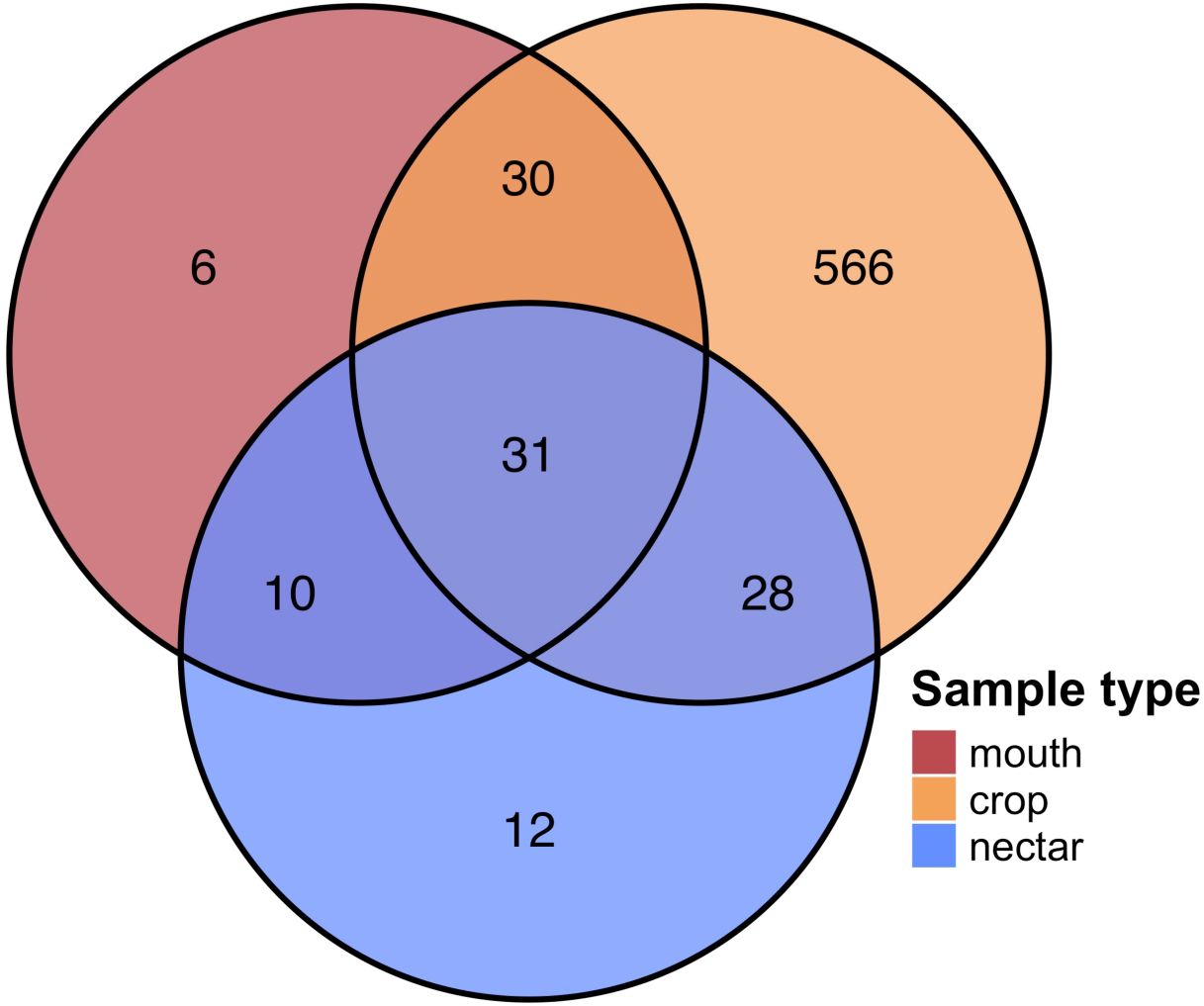
The crop contains more common and constant ASVs_97_ than the mouth and the nectar. Only 9% of 655 winter crop ASVs_97_ were shared with the mouth or nectar samples whereas the mouth and nectar shared about 50% of their 77 and 81 ASVs_97,_ respectively. Venn diagrams are colored by sample type with ASV_97_ counts in bold

**Fig. 3.**
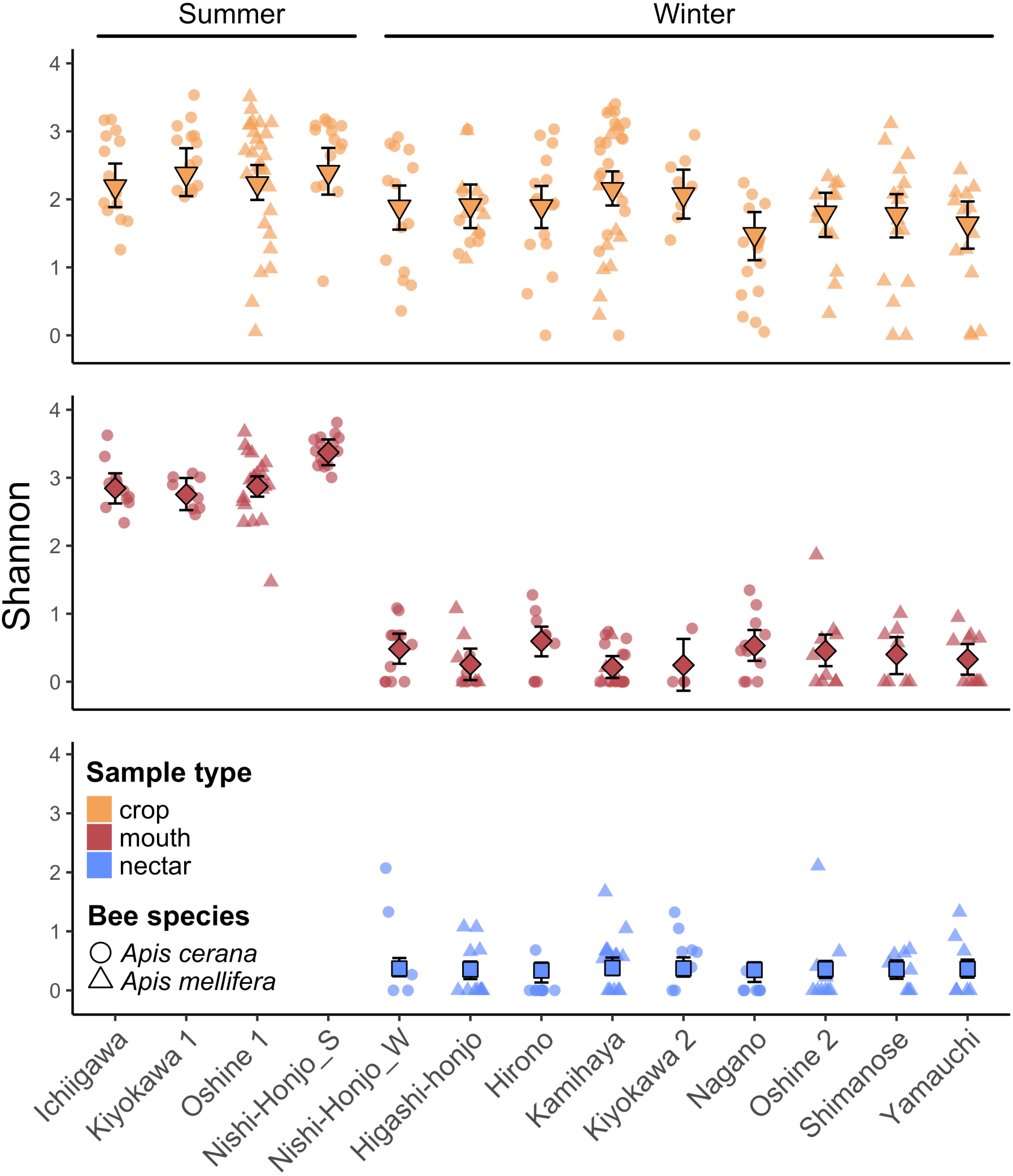
The crop presents more consistent species richness. Species richness (Shannon index) in the crop varied less from summer to winter than the mouth, and winter mouth and nectar samples had similar species richness across sites. Nishi-Honjo_W and Nishi-Honjo_S are the samples collected at the Nishi-Honjo site (see Table S1) in winter and summer, respectively. Each data point represents a single sample collected from the labeled site, colored by sample type, and shaped by host species. Mean diversity is depicted by the large center point and the error bars depict the bootstrapped 95% confidence interval for samples from each site.

Of sample type (crop, mouth, or nectar), bee species (*A. mellifera* or *A. cerana*), and sampling site (Fig. 1a), only sample type was a significant predictor of bacterial ASV_97_ composition (PERMANOVA *R*^2^ = 0.10 *p-value* = 0.001). Specifically, mouth and nectar samples were more similar to each other in bacterial composition than either mouth and crop samples or nectar and crop samples were to each other (Fig. 4; PERMANOVA mouth*–*nectar *R*^2^ = 0.03, *p-value* = 0.002; mouth*–*crop *R*^2^ = 0.11, *p-value* = 0.001; nectar*–*crop *R*^2^ = 0.08, *p-value* = 0.001).

**Fig. 4.**
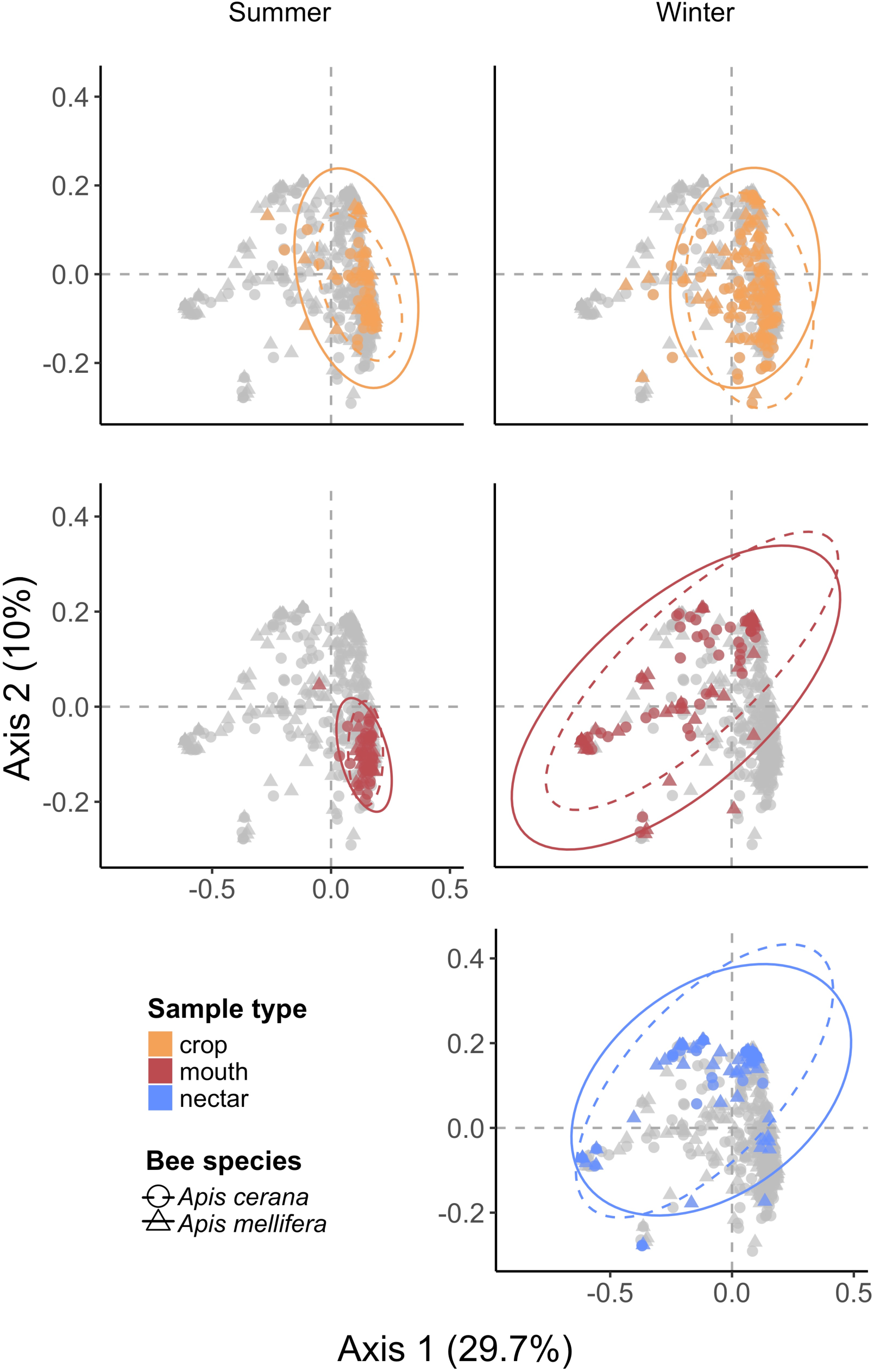
Principal coordinate analysis of all samples presents similar composition across season in the crop but not the mouth. Winter mouth and nectar samples were highly similar (PERMANOVA sample type: *R*^2^ = 0.07, *p-value* = 0.001, season: *R*^2^ = 0.05 *p-value* = 0.001, host species: *R*^2^ = 0.003, *p-value* = 0.154). Each dot represents a sample from an individual bee, colored by sample type. *Apis mellifera* samples are depicted with triangles and the solid ellipse whereas *Apis cerana* samples are depicted with circles and the dashed ellipse. Ellipses mark the 95% confidence interval, and axes are labeled with corresponding percent variation explained

In fact, most ASVs_97_ in the crop were distinct from those in the mouth and the nectar, with only 9% (4-8% in *A. cerana* and 6-9% in *A. mellifera*; Fig. S2) of the crop ASVs_97_ also present in the mouth or the nectar (Fig. 2). Taxa frequently found in the crop, such as *Bombella, Gilliamella, Lactobacillus*, and *Snodgrassella*, were mostly absent in the nectar and only infrequently present in the mouth (Figs. 5 and S3). Furthermore, some ASVs_97_ belonging to *Acinetobacter,* Comamonadaceae, and *Flavobacterium* were consistently found in the crop at all local sites examined, whereas *Acinetobacter* was present in only seven of the nine nectar sites and 11 of the 15 mouth sites, and there were no other taxa that were present in mouth or nectar samples at all sites.

**Fig. 5.**
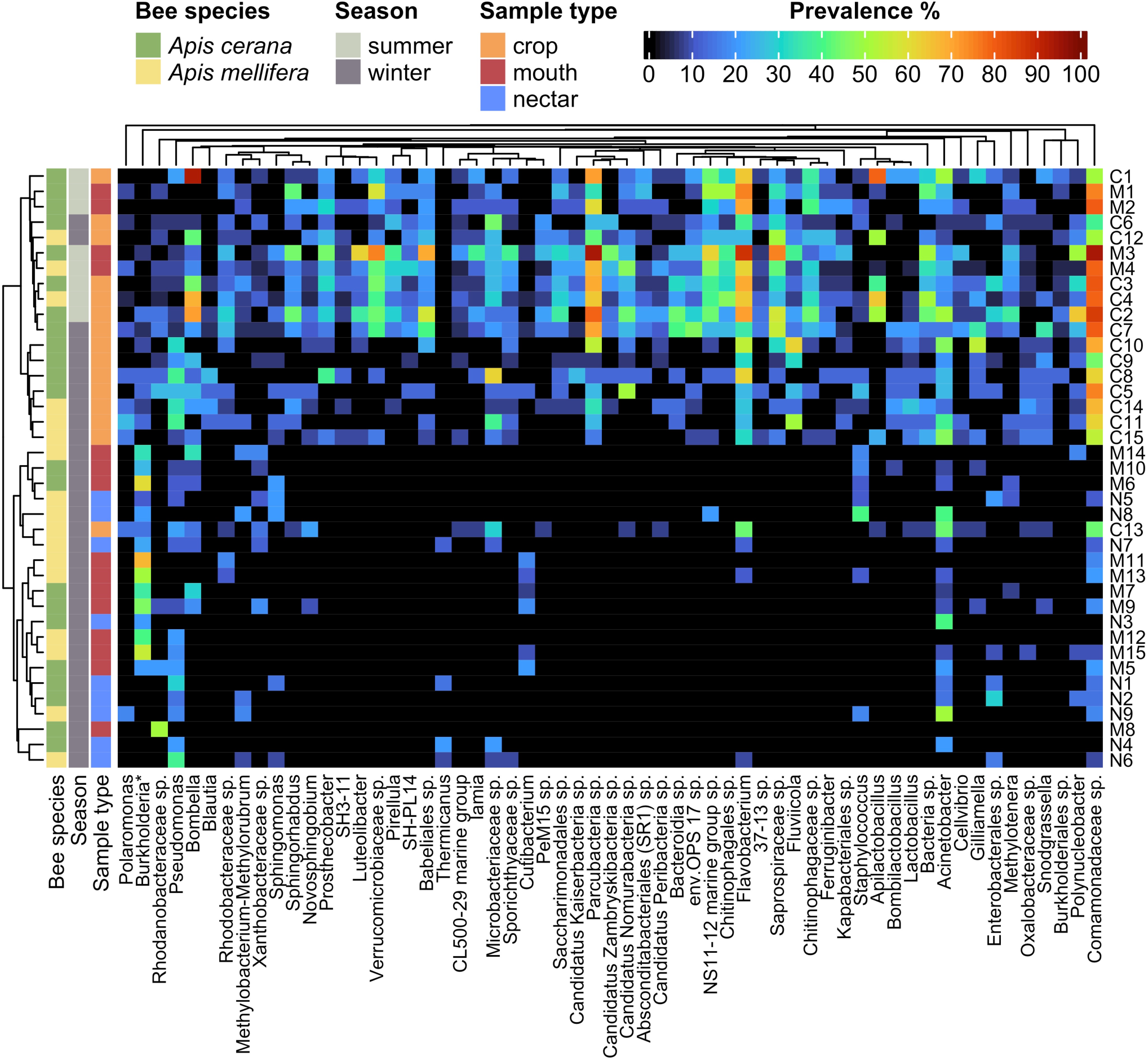
There is a greater prevalence of differentially abundant taxa among crop groups than mouth and nectar groups. Samples were grouped by season, sampling site, host *Apis* species, and sample type as illustrated by colored bars to the left of the heatmap and described in Table S2. For ease of visualization, heatmap columns are differentially abundant taxa that were present in at least 50% of one of the sample groups e.g., at least 50% of all the mouth groups. Burkholderia* denotes the genus Burkholderia-Caballeronia-Paraburkholderia. The full heatmap with individual samples and all differentially abundant taxa can be found in Figure S3. The ASVs_97_ were agglomerated to genus, colored by prevalence within groups, and arranged according to their phylogeny. The y-axis dendrogram depicts the weighted UniFrac distance between samples

These results held true even when we excluded those taxa that are known as the core species of the hindgut bacterial community (*Gilliamella*, *Lactobacillus*, and *Snodgrassella*) from the dataset used for the analysis (Fig. S4).

### Summer–winter difference in crop vs. in mouth

Although ASV_97_ richness was higher in summer than in winter in both the crop and the mouth, the magnitude of this difference was eight times greater in the mouth than in the crop (Figs. 3 and S5). In terms of ASV_97_ composition, most of the crop samples were clustered together regardless of season in both the relative and absolute abundance datasets (relative abundance: winter vs. summer, PERMANOVA *R*^2^ = 0.03, *p-value* = 0.001; absolute abundance: winter vs. summer, PERMANOVA *R*^2^ = 0.04, *p-value* = 0.021), whereas the mouth samples formed two distinct clusters, corresponding to summer and winter (relative abundance: PERMANOVA *R*^2^ = 0.20, *p-value* = 0.001; absolute abundance: PERMANOVA *R*^2^ = 0.31, *p-value* = 0.001; Figs. 4-6a). These two mouth clusters differed in the amount of bacterial composition variation among samples, with winter mouth samples showing higher variation than summer mouth samples (Figs. 4 and 6a) despite no significant difference between them in total bacterial load (Fig. 6b). In contrast to the mouth, the bacterial load in the crop was significantly higher in the summer than in the winter in the *A. mellifera* crop (but not in *A. cerana* crop; Fig. 6b).

**Fig. 6.**
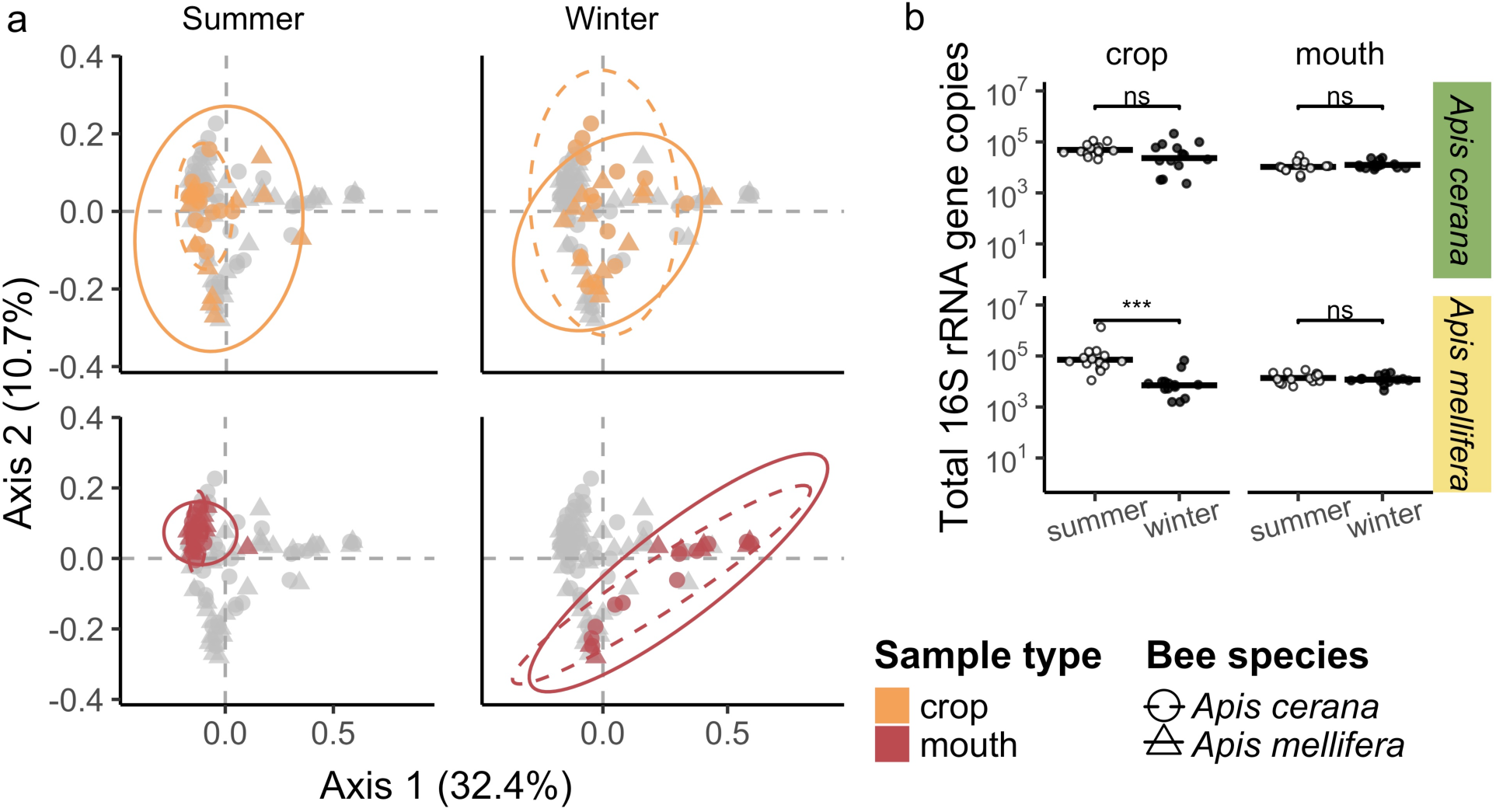
Consideration of absolute abundance estimates in analysis of bacterial composition. (**a**) As in Figure 4, principal coordinate analysis of samples that takes into account absolute total bacterial abundance indicates greater compositional similarity between crop samples than mouth samples across season. Specifically, mouth samples cluster more closely in the summer and are more dispersed in the winter than the crop samples (betadisper distance to median: summer-mouth = 0.21, winter-mouth = 0.40, summer-crop = 0.26, winter-crop = 0.32; betadisper Tukey HSD summer-winter mouth: *p-value* < 0.001, summer-winter crop: *p-value* = 0.008). Overall, sample type and season but not host species are significant predictors of composition (PERMANOVA sample type: *R*^2^ = 0.03, *p-value* = 0.009, season: *R*^2^ = 0.12, *p-value* = 0.001, host species: *R*^2^ = 0.01, *p-value* = 0.214). Each point represents a single sample, is color coded by sample type, and shaped by bee species. Ellipses represent the 95% confidence interval for each bee species. Percent variation explained by each axis is labeled. (**b**) Mouth samples tended to have a more even, lower bacterial load across season than did crop samples. The *Apis mellifera* but not the *Apis cerana* crop samples had a higher bacterial load in the summer than the winter. The mouth maintained the same bacterial load across seasons. Total 16S rRNA gene copies per 10 μl are presented on the log scale, with each point depicting a sample from an individual bee colored by season (summer = white, winter = black). Wilcoxon test significance is labeled with *** = *p-value* < 0.001 and ns = not significant

The CLAM test [25] identified 28% of the ASVs_97_ in the crop as generalists, meaning that they were similarly frequent in winter and summer. In contrast, the CLAM test classified only 2% of the ASVs_97_ in the mouth as generalists, with the majority of others identified as either summer- or winter-associated (76 and 8%, respectively; Fig. 7).

**Fig. 7.**
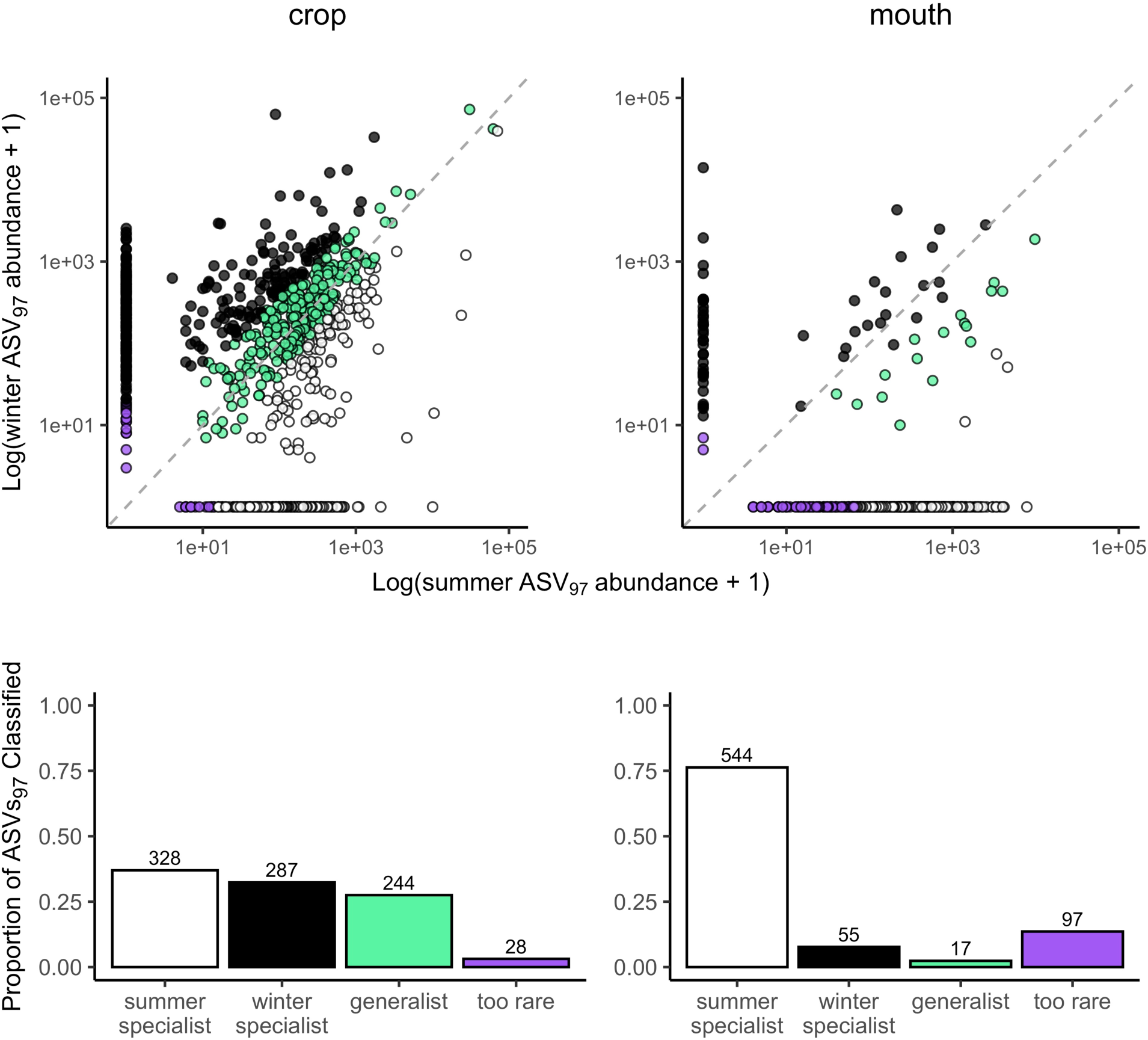
There are bacteria that have high affinity for the crop. Generalist bacteria, i.e., those with no special affinity to one environment or, in this case, season, are greater between the summer and winter crop samples (about 27% of the ASVs_97_) than the corresponding mouth samples. The mouth presents only about 2% generalists, with most ASVs_97_ classified as summer-associated (76%). Scatterplots depict the log abundance of each ASV_97_, with each point representing a single ASV_97_ color coded by its category. ASVs_97_ from both *A. mellifera* and *A. cerana* are included. The identity line is plotted in a dotted grey pattern. The proportion of ASVs_97_ classified in each category is depicted in the corresponding bar plots below the scatterplots, with categories colored as in the scatterplot

### Apilactobacillus kunkeei *strains in the crop*

The multi-gene tree of *Apilactobacillus kunkeei* strains suggested potential season- and *Apis* species-specificity. Specifically, the tree had one clade consisting of only isolates from summer *A. cerana* samples and two other clades consisting of isolates from summer and winter *A. mellifera* samples (Figs. 8 and S6). Those associated with *A. cerana* originated from five bees, three from Nishi-Honjo and two from Ichiigawa. Those associated with *A. mellifera* originated from eight bees: five summer bees from Oshine 1 and three winter bees, one each from Shimanose, Yamauchi, and Kamihaya.

**Fig. 8.**
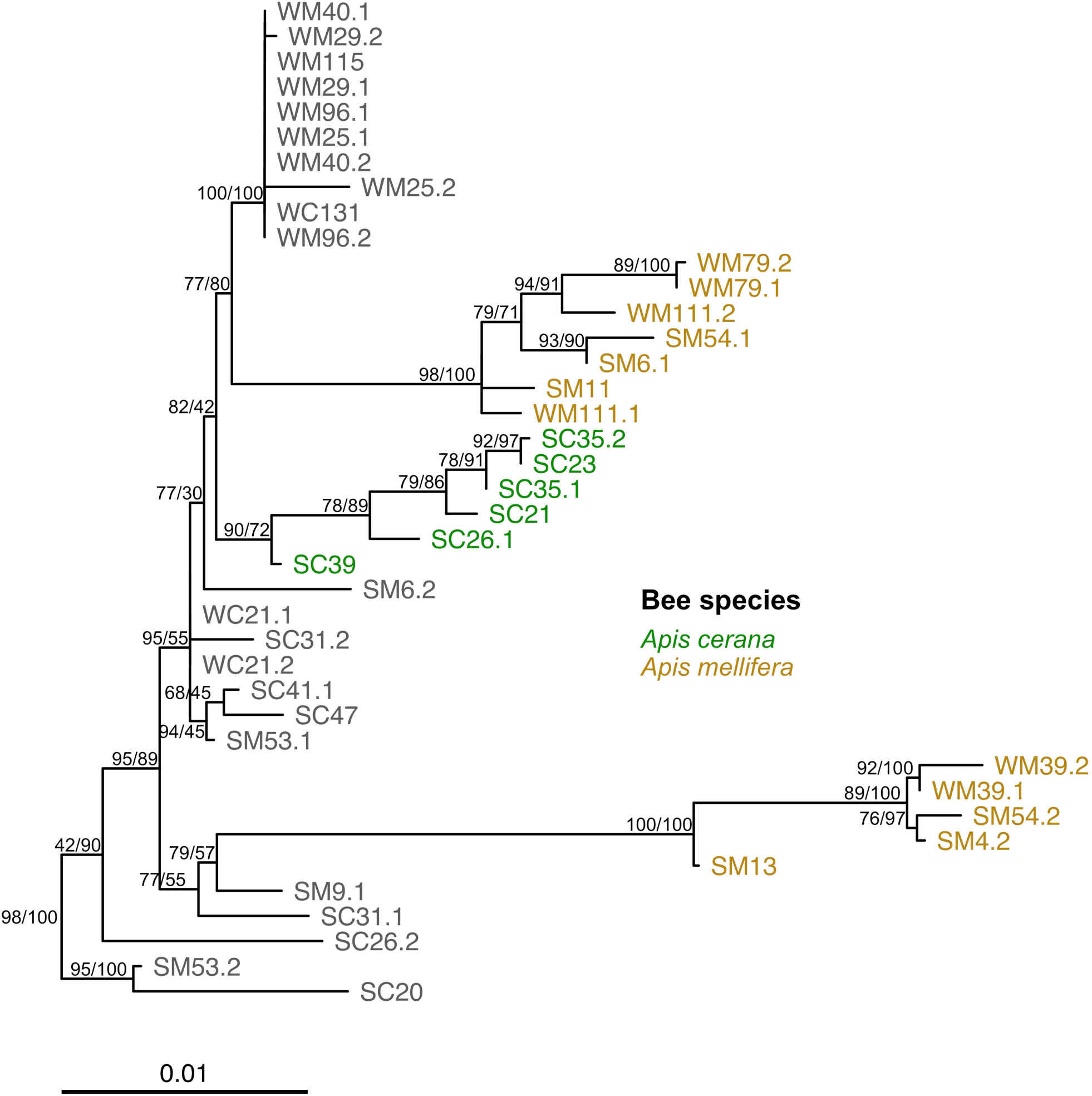
Multi-gene maximum likelihood tree of *Apilactobacillus kunkeei* isolates contains *Apis* species specific clades. The phylogeny is the result of a multi-gene alignment partitioned analysis of 49 *A. kunkeei* isolates and *A. apinorum* Fhon13 (see Tables S3 and S4). In the depicted tree, substitution frequencies are represented by branch length. Isolates are labeled starting with an “S” for summer or “W” for winter, followed by a “C” for *Apis cerana* and an “M” for *Apis mellifera* host species, the host ID number, and lastly the isolate number. Isolates that occurred in both species are colored in grey whereas those that are phylogenetically similar with high statistical support and associated with a single host species are colored by host species. Statistical support was estimated from 1000 Shimodaira-Hasegawa-approximate likelihood ratio test (SH-aLRT) pseudo replicates and ultrafast bootstraps (UFboot). Nodes with an SH-aLRT score greater than 0 are labeled with respective SH-aLRT/UFboot values. *Apilactobacillus apinorum* root is not depicted for ease of visualization. Figure S6 presents the full tree with statistical support on all nodes and *A. apinorum* branch

## Discussion

Based on the conventional assumption that bacterial assemblage in the crop primarily consists of transient microbes acquired via nectar foraging [2, 27, 35, 36], we expected a high degree of bacterial overlap among nectar, mouth, and crop samples. We did find high overlap between the nectar and the mouth in the winter, but to our surprise, most of the crop taxa were clearly distinct from those found in the nectar and the mouth (Figs. 2 and 5) [11]. Several ASVs_97_ of *Acinetobacter* and *Pseudomonas* that were relatively prevalent in both crop and mouth samples in our study have been commonly found in nectar as well [37–39]. However, most of the taxa that were more prevalent in the crop were rare or entirely absent in nectar and mouth samples (Figs. 5, S2 and S3) [11].

In addition, our results indicate that the crop maintained a high degree of taxonomic consistency between winter and summer despite differences in bacterial load in *A. mellifera* crop samples, a pattern that is in sharp contrast to the large compositional changes seen in the mouth between seasons and between bee species (Figs. 3, 4, 6a and 7). Taxa common in the crop in both seasons included Comamonadaceae, *Acinetobacter*, Saprospiraceae, *Flavobacterium*, *Gilliamella*, *Bombella*, *Apilactobacillus*, *Pseudomonas*, *Fluviicola*, *Polynucleobacter*, *Snodgrassella*, and *Microbacteriaceae* species (Fig. 5). *Apilactobacillus* and *Bombella*, previously called *Parasaccharibacter apium* [5, 40], which are among the most commonly reported taxa in the crop of foraging adults, have also been found frequently in food stores, larvae, and queens [2, 4, 5, 27, 36, 41–43]. Although the data we present in this study are correlational, preventing us from establishing causal relationships, we speculate that the crop may serve as a consistent reservoir of these taxa that can influence the health of the adult bees and the larvae that they tend [2, 27].

We also found evidence for potential strain-level specificity among *Apilactobacillus kunkeei* isolates from collected foragers. One clade was made up of summer *A. cerana* isolates from two different sites, and two others contained only *A. mellifera* isolates from both seasons (Figs. 8 and S6). Although a larger sample size than we had in this study as well as full genome sequencing are needed to ascertain the putative strain-level differences, previous studies have shown differences in specific house-keeping genes to suggest strain-level host specificity between *Apis* and *Bombus*, and even among *Apis* species [44, 45]. As in these previous studies, some of our isolates clustered more frequently by host species than by site [46, 47], potentially suggesting host specificity and possibly specific host-symbiont relationships between *Apis* species and *A. kunkeei* strains, even though this possibility remains speculative at this stage.

A few caveats should be considered in interpreting our findings. First, our analysis is mostly based on relative abundance [46], even though we used qPCR to attempt to estimate at least total absolute abundance in crop and mouth samples (Fig. 6b). Absolute abundance of each taxon, which may be more relevant to bacterial effects on the host [12, 46], can differ even when relative abundance is the same. However, we did see similar bacterial composition patterns in our analysis of both relative and absolute abundance (Figs. 4 and 6a). Second, we found that the crop did not consistently have a higher bacterial load than the mouth (Fig. 6b). Although resident populations can be persistent regardless of size, there may be more to the crop community than is suggested by the compositional data. Third, we had only one site, Nishi-Honjo, where bees were sampled in both summer and winter. A greater and more even number of bees at each site in multi-year seasonal sampling would have made our results most robust. Nevertheless, our analysis of several hundred bee and nectar samples all sampled within the same landscape clearly refutes the hypothesis that crop communities simply reflect what is available in the environment.

In conclusion, our results suggest that the composition of crop bacteria may be more deterministic than generally assumed, questioning the notion that crop bacteria are environmentally acquired transients that are unimportant to the bees. Although less well studied than midgut and hindgut microbes, a small but increasing number of recent studies indicate that some crop microbes do alter the health of individuals and hives [6, 48–50]. Our data reinforce the idea that it would help to study how microbial communities are assembled and maintained not just in the midgut and the hindgut, but also the crop, to better understand the relationship between honeybees and their gut microbes.

## Author Contributions

KT, MK, and TF designed the study and collected the samples. MLW did the DNA extraction, sequencing preparation, and bioinformatic analyses. JY and ACH did the quantitative PCR processing and analyses. MLW and LED did the statistical analyses. MLW wrote the first draft of the manuscript. All authors contributed to and approved the manuscript.

## Data Availability

The datasets generated during and/or analyzed during the current study are available in GenBank, https://www.ncbi.nlm.nih.gov (BioProject ID no. PRJNA1076572 and accession no. PP412473-PP412519, PP341993-PP342038, PP382363-PP382409, PP382266-PP382314, PP382315-PP382362), or are included in this published article and its supplementary information files. All custom scripts are available on GitHub at https://github.com/mlwarren20/ume_bee_analysis.

## Acknowledgments

We thank Ryota Nakahaya of Minabe town, Kenichi Hirohata of Tanabe city, and the staff members at the Japanese Apricot Laboratory of the Wakayama Fruit Tree Experiment Station for logistical help; Michiko Futaba, Masaru Horiguchi, Tsutomu Matsuba, Motoki Morikawa, Toshio Nishikawa, Atsuo Okada, Toshiaki Sawa, Yoshinori Taira, Tamotsu Touroku, Ikunori Yamamoto, Toshihiro Yamanouchi, and Shinichi Yamazaki for allowing us to collect bees from their hives and orchards; Junji Takabayashi for laboratory support; Callie Chappell for graphical abstract help; and Taro Maeda, Junichi Takahashi, and the members of the community ecology group at Stanford University for discussion. We also thank Josh Neufeld, Irene Newton, Julie Olson, and five anonymous reviewers for comments on early versions of this manuscript. This work was supported by the Office of International Affairs and the Freeman Spogli Institute for International Studies Japan Fund of Stanford University.

## References

1. Kwong WK, Moran NA (2016) Gut microbial communities of social bees. Nat Rev Microbiol 14: 374–384. doi: 10.1038/nrmicro.2016.43

2. Anderson KE, Ricigliano VA, Copeland DC, Mott BM, Maes P (2023) Social interaction is unnecessary for hindgut microbiome transmission in honey bees: the effect of diet and social exposure on tissue-specific microbiome assembly. Microb Ecol 85: 1498–1513. doi: 10.1007/s00248-022-02025-5

3. Winston ML (1987) The biology of the honey bee. harvard university press

4. Tamarit D, Ellegaard KM, Wikander J, Olofsson T, Vásquez A, Andersson SGE (2015) Functionally structured genomes in *Lactobacillus kunkeei* colonizing the honey crop and food products of honeybees and stingless bees. Genome Biol Evol 7: 1455–1473. doi: 10.1093/gbe/evv079

5. Corby-Harris V, Snyder LA, Schwan MR, Maes P, McFrederick QS, Anderson KE (2014) Origin and effect of Alpha 2.2 Acetobacteraceae in honey bee larvae and description of *Parasaccharibacter apium* gen. nov., sp. nov. Appl Environ Microbiol 80: 7460–7472. doi: 10.1128/AEM.02043-14

6. Arredondo D, Castelli L, Porrini MP, Garrido PM, Eguaras MJ, Zunino P, Antúnez K (2018) *Lactobacillus kunkeei* strains decreased the infection by honey bee pathogens *Paenibacillus larvae* and *Nosema ceranae*. Benef Microbes 9: 279–290. doi: 10.3920/BM2017.0075

7. Kwong WK, Mancenido AL, Moran NA (2017) Immune system stimulation by the native gut microbiota of honey bees. Roy Soc Open Sci 4: 170003.

8. Zheng H, Powell JE, Steele MI, Dietrich C, Moran NA (2017) Honeybee gut microbiota promotes host weight gain via bacterial metabolism and hormonal signaling. Proc Natl Acad Sci USA 114: 4775–4780. doi: 10.1073/pnas.1701819114

9. Hart AG, Ratnieks FLW (2001) Why do honey-bee (*Apis millifera*) foragers transfer nectar to several receivers? Information improvement through multiple sampling in a biological system. Behav Ecol Sociobiol 49: 244–250. doi: 10.1007/s002650000306

10. Decker LE, San Juan PA, Warren ML, Duckworth CE, Gao C, Fukami T (2022) Higher variability in fungi compared to bacteria in the foraging honey bee gut. Microb Ecol 85: 330–334. doi: 10.1007/s00248-021-01922-5

11. Tiusanen M, Becker-Scarpitta A, Wirta H (2024) Distinct Communities and Differing Dispersal Routes in Bacteria and Fungi of Honey Bees, Honey, and Flowers. Microb Ecol 87: 100. doi: 10.1007/s00248-024-02413-z

12. Kešnerová L, Emery O, Troilo M, Liberti J, Erkosar B, Engel P (2020) Gut microbiota structure differs between honeybees in winter and summer. ISME J 14: 801–814. doi: 10.1038/s41396-019-0568-8

13. Maeda T, Hiraiwa MK, Shimomura Y, Oe T (2023) Weather conditions affect pollinator activity, fruit set rate, and yield in Japanese apricot. Sci Hortic 307: 111522. doi: 10.1016/j.scienta.2022.111522

14. R Core Team (2023) R: a language and environment for statistical computing. In: Computing, RffS (ed.), Vienna, Austria.

15. McMurdie PJ, Holmes S (2013) phyloseq: An R package for reproducible interactive analysis and graphics of microbiome census data. PLoS One 8: e61217. doi: 10.1371/journal.pone.0061217

16. Oksanen J, Blanchet FG, Friendly M, Kindt R, Legendre P, McGlinn D, Minchin PR, O’Hara RB, Simpson GL, Solymos P, Stevens MHH, Szoecs E, Wagner H (2019) vegan: Community ecology package. R package version 2.5–6.

17. Xu S, Zhan L, Tang W, Wang Q, Dai Z, Zhou L, Feng T, Chen M, Wu T, Hu E (2023) MicrobiotaProcess: A comprehensive R package for deep mining microbiome. Innovation 4.

18. Chen H (2022) VennDiagram: generate high-resolution Venn and Euler plots. R package version 1.7.3.

19. Love MI, Huber W, Anders S (2014) Moderated estimation of fold change and dispersion for RNA-seq data with DESeq2. Genome Biol 15: 550. doi: 10.1186/s13059-014-0550-8

20. Gu Z, Eils R, Schlesner M (2016) Complex heatmaps reveal patterns and correlations in multidimensional genomic data. Bioinformatics 32: 2847–2849. doi: 10.1093/bioinformatics/btw313

21. Schliep K (2011) phangorn: phylogenetic analysis in R. Bioinformatics 27: 592e593.

22. Hamady M, Lozupone C, Knight R (2010) Fast UniFrac: facilitating high-throughput phylogenetic analyses of microbial communities including analysis of pyrosequencing and PhyloChip data. ISME J 4: 17–27.

23. Bates D, Mächler M, Bolker B, Walker S (2015) Fitting linear mixed-effects models using lme4. J Stat Softw 67: 1–48. doi: 10.18637/jss.v067.i01

24. Lahti L, Shetty S (2017) microbiome R package.

25. Chazdon RL, Chao A, Colwell RK, Lin S-Y, Norden N, Letcher SG, Clark DB, Finegan B, Arroyo JP (2011) A novel statistical method for classifying habitat generalists and specialists. Ecology 92: 1332–1343. doi: 10.1890/10-1345.1

26. Ogier J-C, Pagès S, Galan M, Barret M, Gaudriault S (2019) *rpoB*, a promising marker for analyzing the diversity of bacterial communities by amplicon sequencing. BMC Microbiol 19: 171. doi: 10.1186/s12866-019-1546-z

27. Dyrhage K, Garcia-Montaner A, Tamarit D, Seeger C, Näslund K, Olofsson TC, Vasquez A, Webster MT, Andersson SG (2022) Genome evolution of a symbiont population for pathogen defense in honeybees. Genome Biol Evol 14: evac153.

28. Katoh K, Standley DM (2013) MAFFT Multiple sequence alignment software version 7: improvements in performance and usability. Mol Biol Evol 30: 772–780. doi: 10.1093/molbev/mst010

29. Capella-Gutiérrez S, Silla-Martínez JM, Gabaldón T (2009) trimAl: a tool for automated alignment trimming in large-scale phylogenetic analyses. Bioinformatics 25: 1972–1973.

30. Minh BQ, Schmidt HA, Chernomor O, Schrempf D, Woodhams MD, von Haeseler A, Lanfear R (2020) IQ-TREE 2: new models and efficient methods for phylogenetic inference in the genomic era. Mol Biol Evol 37: 1530–1534. doi: 10.1093/molbev/msaa015

31. Anisimova M, Gil M, Dufayard J-F, Dessimoz C, Gascuel O (2011) Survey of branch support methods demonstrates accuracy, power, and robustness of fast likelihood-based approximation schemes. Syst Biol 60: 685–699. doi: 10.1093/sysbio/syr041

32. Hoang DT, Chernomor O, von Haeseler A, Minh BQ, Vinh LS (2017) UFBoot2: improving the ultrafast bootstrap approximation. Mol Biol Evol 35: 518–522. doi: 10.1093/molbev/msx281

33. Kalyaanamoorthy S, Minh BQ, Wong TKF, von Haeseler A, Jermiin LS (2017) ModelFinder: fast model selection for accurate phylogenetic estimates. Nat Methods 14: 587–589. doi: 10.1038/nmeth.4285

34. Mai U, Mirarab S (2018) TreeShrink: fast and accurate detection of outlier long branches in collections of phylogenetic trees. BMC Genomics 19: 23–40.

35. Anderson KE, Maes P (2022) Social microbiota and social gland gene expression of worker honey bees by age and climate. Sci Rep 12: 10690. doi: 10.1038/s41598-022-14442-0

36. Anderson KE, Sheehan TH, Mott BM, Maes P, Snyder L, Schwan MR, Walton A, Jones BM, Corby-Harris V (2013) Microbial ecology of the hive and pollination landscape: bacterial associates from floral nectar, the alimentary tract and stored food of honey bees (*Apis mellifera*). PLoS One 8: e83125.

37. Alvarez-Perez S, Baker LJ, Morris MM, Tsuji K, Sanchez VA, Fukami T, Vannette RL, Lievens B, Hendry TA (2021) *Acinetobacter pollinis* sp. nov., *Acinetobacter baretiae* sp. nov. and *Acinetobacter rathckeae* sp. nov., isolated from floral nectar and honey bees. Int J Syst Evol Microbiol 71: 004783.

38. Fridman S, Izhaki I, Gerchman Y, Halpern M (2012) Bacterial communities in floral nectar. Environ Microbiol Rep 4: 97–104. doi: 10.1111/j.1758-2229.2011.00309.x

39. Morris MM, Frixione NJ, Burkert AC, Dinsdale EA, Vannette RL (2020) Microbial abundance, composition, and function in nectar are shaped by flower visitor identity. FEMS Microbiol Ecol 96: fiaa003. doi: 10.1093/femsec/fiaa003

40. Smith EA, Anderson KE, Corby-Harris V, McFrederick QS, Parish AJ, Rice DW, Newton ILG (2021) Reclassification of seven honey bee symbiont strains as *Bombella apis*. Int J Syst Evol Microbiol 71: 004950. doi: 10.1099/ijsem.0.004950

41. Vásquez A, Forsgren E, Fries I, Paxton RJ, Flaberg E, Szekely L, Olofsson TC (2012) Symbionts as major modulators of insect health: lactic acid bacteria and honeybees. PLoS One 7: e33188. doi: 10.1371/journal.pone.0033188

42. Corby-Harris V, Maes P, Anderson KE (2014) The bacterial communities associated with honey bee (*Apis mellifera*) foragers. PLoS One 9: e95056.

43. Rangberg A, Mathiesen G, Amdam GV, Diep DB (2015) The paratransgenic potential of *Lactobacillus kunkeei* in the honey bee *Apis mellifera*. Benef Microbes 6: 513–523. doi: 10.3920/BM2014.0115

44. Kwong WK, Engel P, Koch H, Moran NA (2014) Genomics and host specialization of honey bee and bumble bee gut symbionts. Proc Natl Acad Sci USA 111: 11509–11514. doi: 10.1073/pnas.1405838111

45. Kwong WK, Medina LA, Koch H, Sing K-W, Soh EJY, Ascher JS, Jaffé R, Moran NA (2017) Dynamic microbiome evolution in social bees. Sci Adv 3: e1600513. doi: doi:10.1126/sciadv.1600513

46. Almeida EL, Ribiere C, Frei W, Kenny D, Coffey MF, O’Toole PW (2023) Geographical and seasonal analysis of the honeybee microbiome. Microb Ecol 85: 765–778. doi: 10.1007/s00248-022-01986-x

47. Ellegaard KM, Suenami S, Miyazaki R, Engel P (2020) Vast differences in strain-level diversity in the gut microbiota of two closely related honey bee species. Curr Biol 30: 2520–2531.e2527. doi: 10.1016/j.cub.2020.04.070

48. Chege M, Kinyua J, Paredes JC (2023) *Lactobacillus kunkeei* impacts the health of honey bees, *Apis mellifera* scutellata, and protects the bees against the opportunistic pathogen *Serratia marcescens*. Int J Trop Insect Sci 43: 1947–1955. doi: 10.1007/s42690-023-01103-6

49. Corby-Harris V, Snyder L, Meador CAD, Naldo R, Mott B, Anderson KE (2016) *Parasaccharibacter apium*, gen. nov., sp. nov., improves honey bee (Hymenoptera: Apidae) resistance to Nosema. J Econ Entomol 109: 537–543. doi: 10.1093/jee/tow012

50. Miller DL, Smith EA, Newton ILG (2021) A bacterial symbiont protects honey bees from fungal disease. mBio 12: e00503–00521. doi: 10.1128/mBio.00503-21

