## Supplemental Figure 1 for "Bacteria in honeybee crops are decoupled from those in floral nectar and bee mouths"

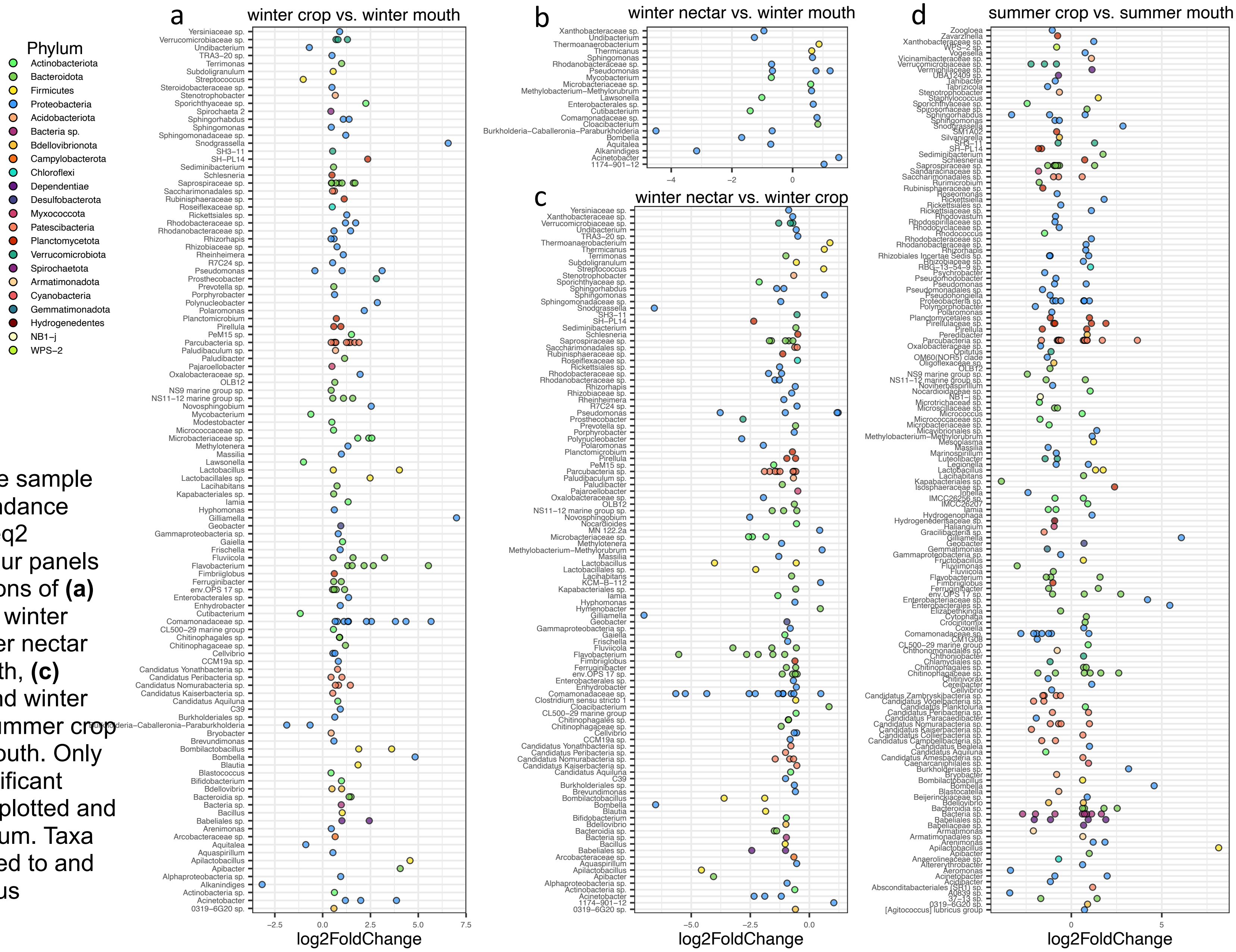


**Fig S1**. Pairwise sample differential abundance based on DESeq2 analysis. The four panels show comparisons of **(a)** winter crop and winter mouth, **(b)** winter nectar and winter mouth, **(c)** winter nectar and winter crop, and **(d)** summer crop and summer mouth. Only statistically significant differences are plotted and colored by Phylum. Taxa are agglomerated to and labeled by Genus
