## Supplemental Figure 2 for "Bacteria in honeybee crops are decoupled from those in floral nectar and bee mouths"

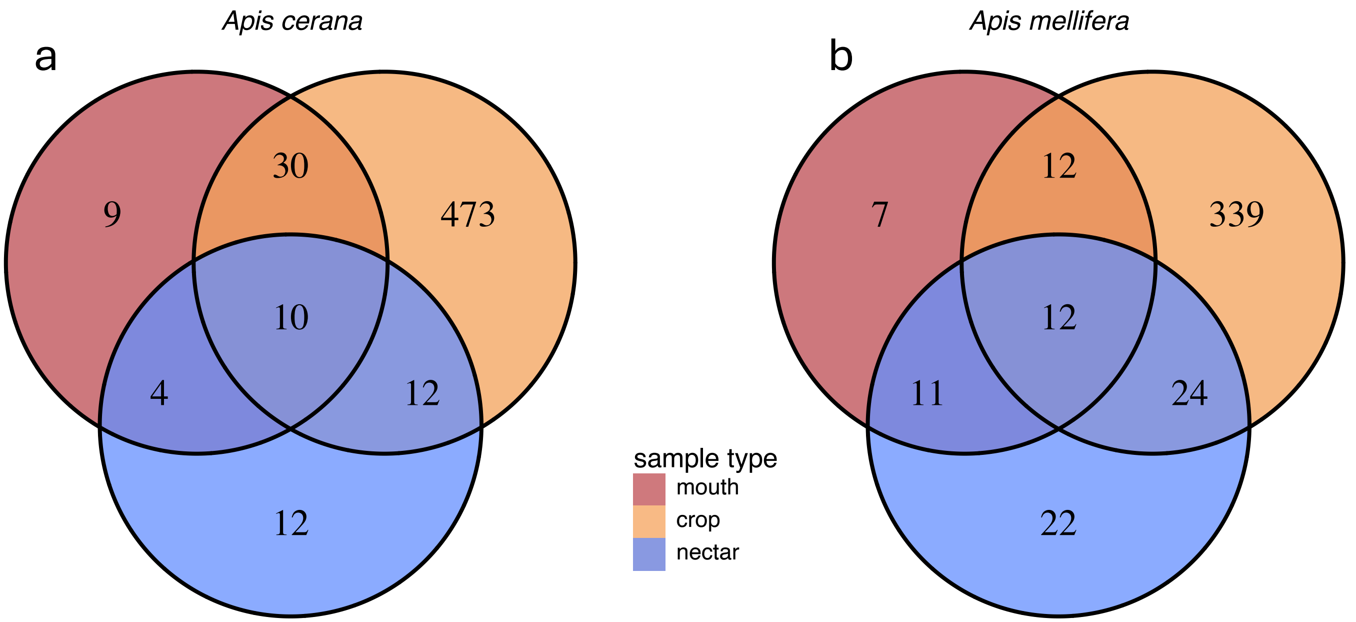


**Fig S2.** Samples collected from *Apis cerana* and *Apis mellifera* sites show the same overall pattern of more common and constant $ASVs_{97}$ in the crop than the mouth and the nectar. (**a**) At *Apis cerana* sites, only 8% and 4% of the winter crop $ASVs_{97}$ were shared with the mouth and nectar samples, respectively, while the mouth and nectar shared about 30% of their $ASVs_{97}$. (**b**) Similarly, at *Apis mellifera* sites, 6 and 9% of the crop $ASVs_{97}$ were shared with the mouth and nectar samples, respectively. Half of the mouth $ASVs_{97}$ were also present in the nectar, while only about 30% of the nectar $ASVs_{97}$ were found in the mouth. Venn diagrams are colored by sample type with $ASVs_{97}$ counts in bold as in Figure 2
