## Supplemental Figure 3 for "Bacteria in honeybee crops are decoupled from those in floral nectar and bee mouths"

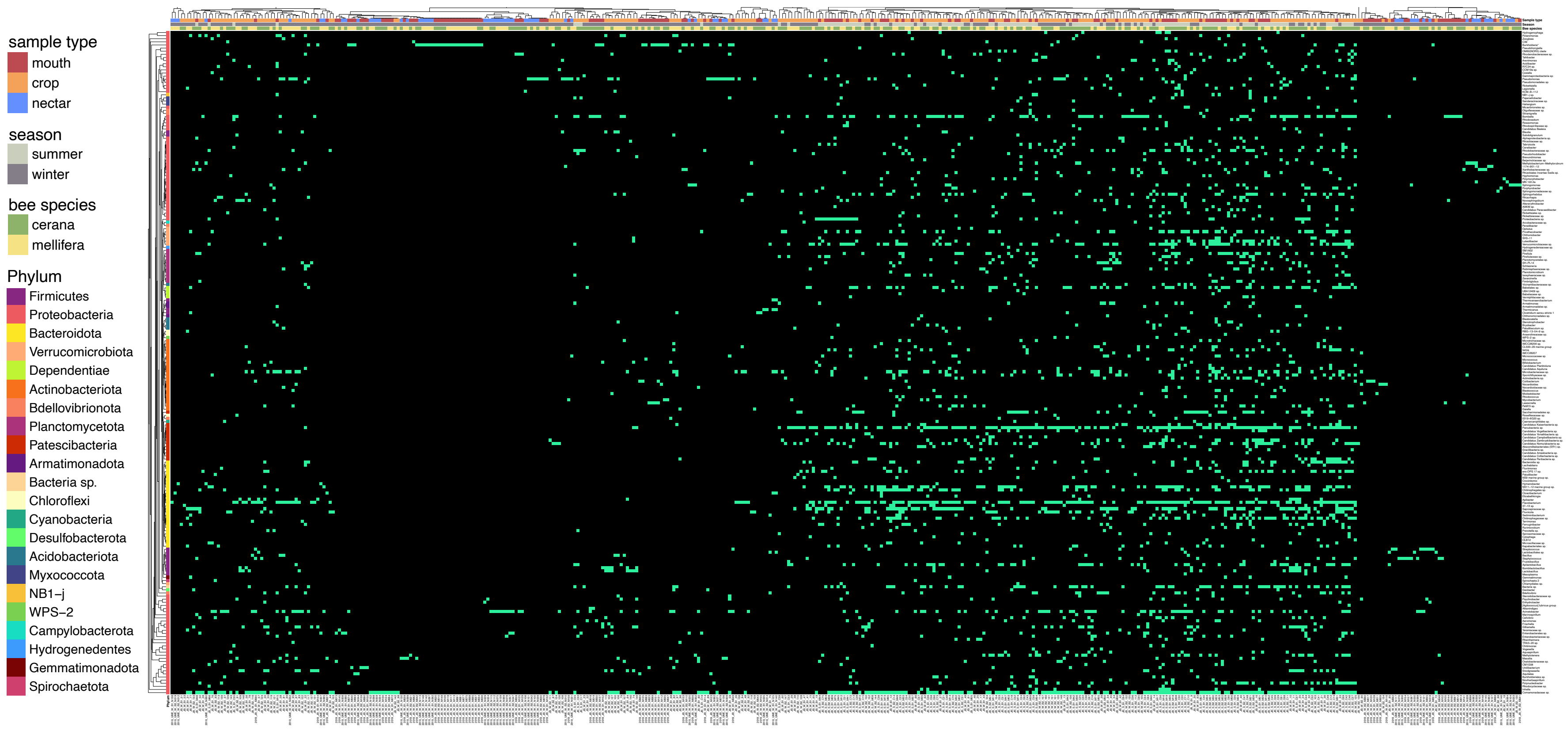


**Fig S3.** Heatmap as in Fig. 6 except that samples are ungrouped and the heatmap represents presence (in green) and absence (in black) of the taxa in each sample. Also, all differentially abundant taxa (see Supplementary Figure 1), agglomerated to genus, are depicted.
