## Supplemental Figure 4 for "Bacteria in honeybee crops are decoupled from those in floral nectar and bee mouths"

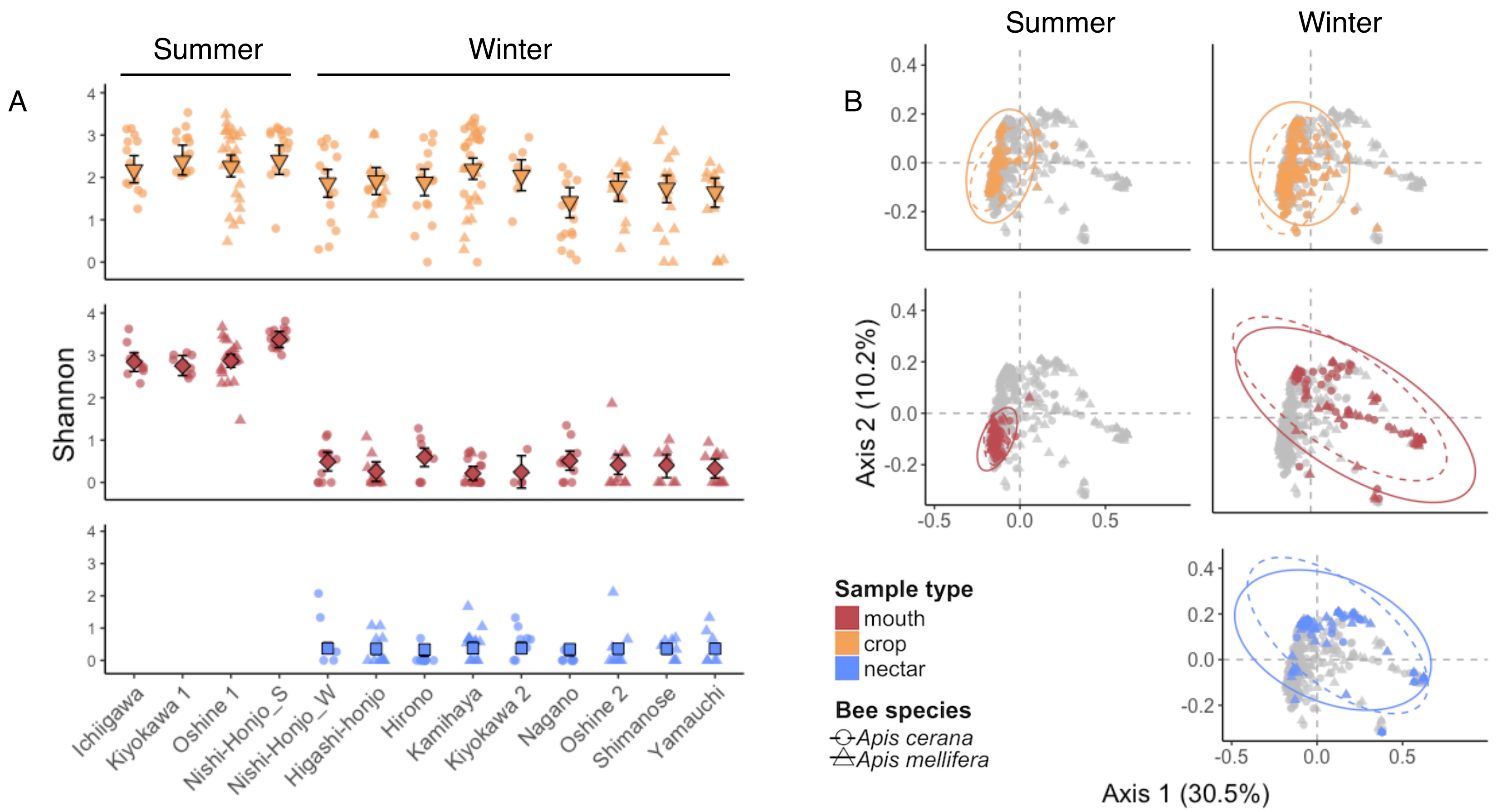


**Fig S4.** Exclusion of genera commonly found in the hindgut, specifically *Gilliamella, Snodgrassella,* and *Lactobacillus*, does not change the (**a**) alpha or (**b**) beta diversity trends of the bacterial communities. The crop bacterial alpha diversity and composition remain more constant across season than those of the mouth, while the mouth and nectar present very similar alpha and beta diversity in the winter.
