## Supplemental Figure 5 for "Bacteria in honeybee crops are decoupled from those in floral nectar and bee mouths"

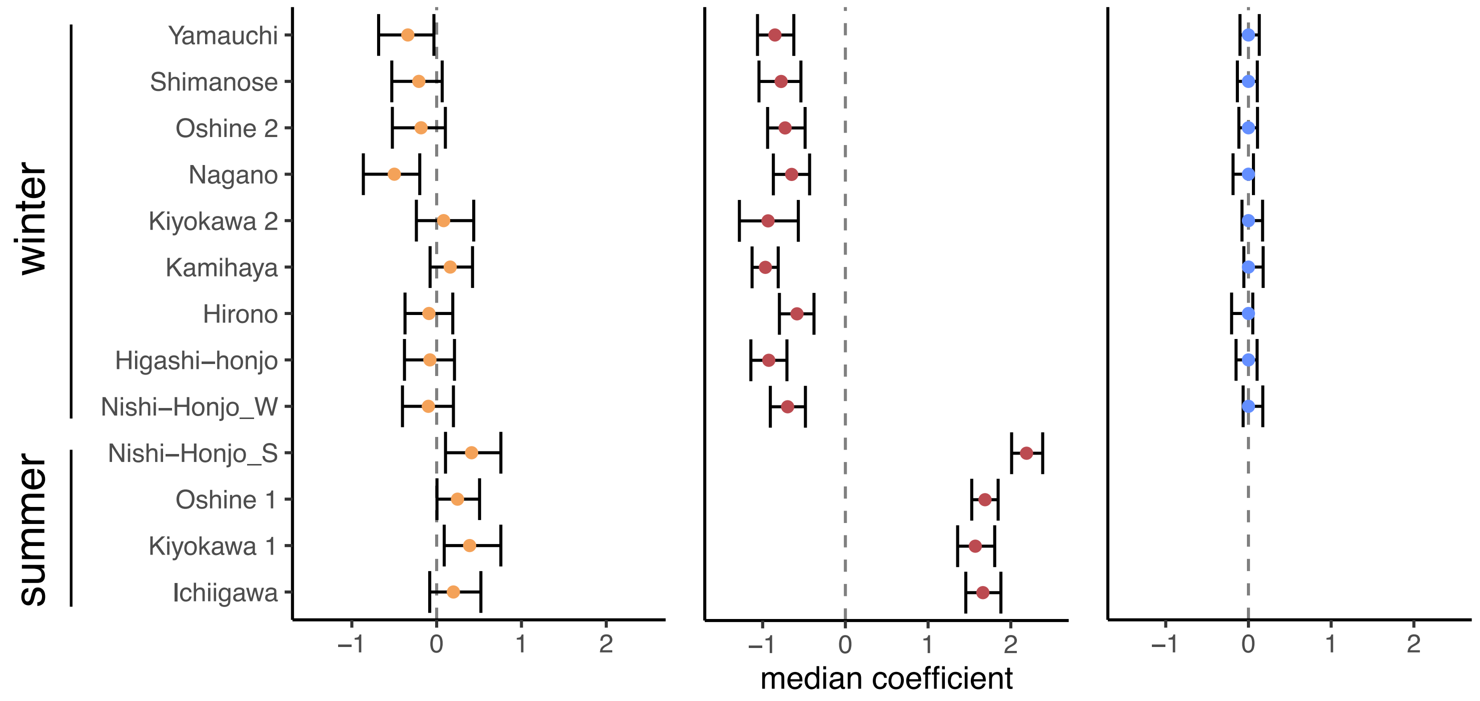


**Fig S5.** Median coefficients with error bars representing the bootstrapped 95% confidence interval of linear mixed effects model of Shannon diversity with the random effect of site.
