## Supplemental Figure 6 for "Bacteria in honeybee crops are decoupled from those in floral nectar and bee mouths"

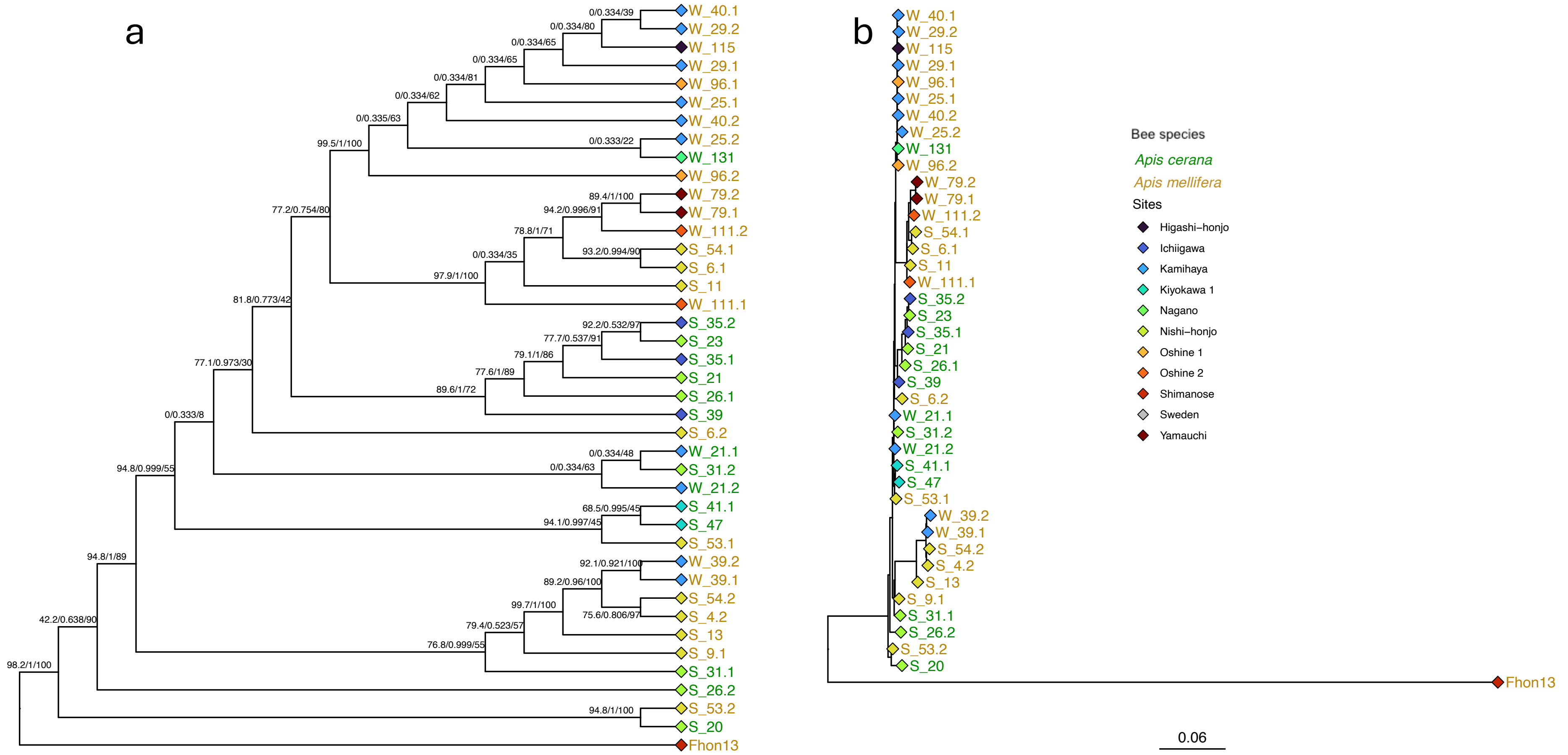


**Fig S6.** (**a**) Cladogram of phylogeny as in Fig. 8 including the outgroup, *Apilactobacillus apinorum* Fhon13, and statistical support from 1000 SH-aLRT pseudo replicates (%), a Bayesian-like transformation of aLRT (aBayes), and 1000 ultrafast bootstraps (%) in that order. (**b**) Multi-gene phylogeny including Fhon13. Branch length represents substitution frequency. Nodes are colored by collection site; sample labels start with an “S” for summer or a “W” for winter samples followed by the host bee ID and the isolate number and are colored by host species.
