## Supplemental Text 1 for "Bacteria in honeybee crops are decoupled from those in floral nectar and bee mouths"

**Text S1**. Microbial DNA analysis

*Microbial DNA extraction*

In our laboratory at Stanford University, the Qiagen DNeasy Blood and Tissue kit and Qiagen Genomic DNA Buffer set (Qiagen, Germantown, MD, USA) were used to extract DNA from all the samples. The extraction followed the gram-positive bacteria protocol along with the following modifications. For each of the crop, mouth, and nectar samples collected, 10 µL of the thawed sample was combined with 2 µL of zymolyase enzyme (MP Biomedicals, Burlingame, CA, USA) within a microcentrifuge tube. Two DNA extraction negative controls, consisting of only the 2 µL of zymolyase enzyme, were included for every 90 samples for a total of 13 DNA extraction negative controls. These mixtures were incubated at 30°C for 30 minutes. Once the incubation period was complete, 10 µL of a 500:1 G2 (from the Qiagen Genomic DNA Buffer set) buffer to RNAse A (Qiagen) mixture was added to each microcentrifuge tube. 20 µL of ProteinaseK (Qiagen) and 180 µL of Buffer ATL (Qiagen) were added to each sample and vortexed to mix. The samples were then incubated at 56°C overnight. The next morning the samples were mixed and centrifuged to collect any condensation on the caps, and 410 µL of AL-Ethanol (Qiagen) mixture were added to each sample and shaken vigorously to mix. The rest of the protocol was followed as stated in the DNeasy Blood & Tissue Handbook (2018) with the only exception being that the final elution of DNA was done with 50 µL of Buffer AE (Qiagen) rather than 200 µL to increase the final DNA concentration in the eluate. The final extracted DNA from each sample was stored in sterile microcentrifuge tubes at 4°C until library preparation.

*Bacterial amplicon sequencing*

The highly variable (V4) region of the 16S rRNA gene was amplified using primers 515F (5´-TCGTCGGCAGCGTCAGATGTGTATAAGAGACAGGTGYCAGCMGCCGCGGTAA-3´) and 806R (5´-GTCTCGTGGGCTCGGAGATGTGTATAAGAGACAGGGACTACNVGGGTWTCTAAT-3´) [1]. Both primers included an Illumina (San Diego, CA, USA) adapter on the 5´ ends used to attach unique Nextera XT indexes for sample identification. Each 10 µL first step PCR amplification consisted of 3.2 µL of PCR-grade water, 5 µL of MyTaq HS Red Mix (Bioline, Taunton, MA, USA), 0.4 µL each of the forward and reverse primers, and 1 µL of the extracted DNA sample. Two negative control samples, where the reaction did not include any input DNA, per every 90 experimental samples, were included for a total of 13 first step PCR negative controls. PCR conditions were 95ºC for 2 minutes, followed by 35 cycles of 95ºC for 20 seconds, 50ºC for 20 seconds, 72ºC for 50 seconds, and a final extension at 72ºC for 10 minutes with a final hold at 4ºC. Amplification was confirmed by running samples on an agarose gel, and the resulting amplicons were purified using Sera-Mag SpeedBeads (Sigma-Aldrich, St. Louis, MO, USA). The second step 10 µL PCR amplification consisted of 3.2 µL of PCR-grade water, 5 µL of MyTaq HS Red Mix (Bioline), 0.4 µL each of the combined Nextera XT index primers 1 and 2, and 1 µL of the first step PCR product. Two negative control samples, where the reaction did not include any of the first step PCR product, per every 90 experimental samples, were included for a total of 13 second step PCR negative controls. PCR conditions for this second step were 95ºC for 2 minutes, followed by 8 cycles of 95ºC for 20 seconds, 50ºC for 20 seconds, 72ºC for 50 seconds, and a final extension at 72ºC for 10 minutes with a final hold at 4ºC. Amplification was confirmed by running samples on an agarose gel, the barcoded amplicons were purified once again using Sera-mag SpeedBeads (Sigma-Aldrich), and the resulting clean product was pooled evenly across samples. The pooled samples were combined to a concentration of 4 nM using a Qubit 4 Fluorometer (Thermo Fisher, Waltham, MA, USA), and then the entire amplicon libraries were pooled together. The final DNA concentration and quality of the pooled amplicon libraries were quantified with a Fragment Analyzer (Agilent, Santa Clara, CA, USA), and then sequenced on a MiSeq (Illumina) using a 2x300 cycle sequencing kit with a 15% PhiX spike-in at the Stanford Genomic Sequencing Service Center.

*Quantitative real-time PCR (qPCR) of 16S rRNA gene*

A subset of the crop and mouth samples was selected for 16S rRNA gene quantification. These 112 samples were made up of mouth and crop samples from 28 *Apis mellifera* and 28 *A. cerana* bees, 14 each from the summer and winter sites. To ensure we captured differences in bacterial load in Nishi-Honjo, the one site where we did the crop and mouth sampling in both seasons, we included all samples from this site in this analysis. The remaining *A. cerana* and *A. mellifera* samples were randomly selected from the other sites. The total number of bacterial 16S rRNA genes was quantified in the extracted DNA samples. Targeted genes were amplified using the CFX96^TM^ Real-Time System (C1000TM Thermal Cycler, Bio-Rad, Hercules, CA, USA). All reactions were performed using SsoFast™ EvaGreen® Supermix (Bio-Rad) according to the manufacturer’s instructions and in triplicate using 10 μL of SsoFast EvaGreen Supermix, 0.6 μL of each primer, 2 μL DNA samples, and nuclease-free water to a final volume of 20 μL. 16S rRNA genes were amplified using the primer sets 1055YF(5’-ATGGYTGTCGTCAGCT-3’) and 1392R (5’-ACGGGCGGTGTGTAC-3’), with the PCR conditions as follows: 2 minutes at 50°C and 10 minutes at 95°C, followed by 40 cycles of 15 seconds at 95°C and 1 minute at 58°C [2]. The detection limit (1.6 x 10^2^ gene copies per reaction) for quantitative real-time PCR was determined using 55 standard curves for 16S rRNA genes that were constructed using the serial 10-fold dilutions from 10^-1^ to 10^-7^ of known concentrations of synthesized oligonucleotides (Integrated DNA Technologies, Inc., Coralville, IA, USA). The gene copy number was determined from CT values equal to or higher than the negative controls were considered as below the detection limit.

*16S rRNA gene amplicon-sequencing data processing*

The raw amplicon sequences were trimmed using the Cutadapt software [3]. The Divisive Amplicon Denoising Algorithm 2 (DADA2) pipeline [4] was used to merge paired-end sequences, quality filter these sequences, remove chimeric reads, and cluster sequences into amplicon sequence variants (ASVs). Taxonomy assignments for each ASV were assigned using the SILVA version 138.1 database [5]. ASVs present in the DNA extraction and first and second step PCR negative controls as well as those identified as chloroplast or mitochondria were removed from all samples in the dataset. Any phyla less than 1% abundant among all samples were trimmed [6]. This preprocessing resulted in several samples containing no sequence reads. Consequently, these samples were excluded from further analysis, resulting in a total of 218 crop samples, 161 mouth samples, and 82 nectar samples used for downstream analysis. To minimize grouping ASVs from the same genome into separate clusters [7], ASVs were grouped into ASVs_97_ sequences, as previously described [8], with a 97% similarity threshold using the DECIPHER and speedyseq R packages [9, 10]. Counts for all samples were transformed to relative abundance by dividing the amount of each ASV_97_ in each individual sample by the total number of reads in the sample. To convert the counts to absolute abundance, for the subset of samples with qPCR data, we multiplied the total number of 16s rRNA gene copies in each sample by the proportion of each ASV_97_ in said sample.

**References**

1.	Caporaso JG, Lauber CL, Walters WA, Berg-Lyons D, Huntley J, Fierer N, Owens SM, Betley J, Fraser L, Bauer M, Gormley N, Gilbert JA, Smith G, Knight R (2012) Ultra-high-throughput microbial community analysis on the Illumina HiSeq and MiSeq platforms. ISME J 6: 1621-1624. doi: 10.1038/ismej.2012.8

2.	Ritalahti KM, Amos BK, Sung Y, Wu Q, Koenigsberg SS, Löffler FE (2006) Quantitative PCR targeting 16S rRNA and reductive dehalogenase genes simultaneously monitors multiple *Dehalococcoides* strains. Appl Environ Microbiol 72: 2765-2774.

3.	Martin M (2011) Cutadapt removes adapter sequences from high-throughput sequencing reads. EMBnet J 17: 10-12. doi: 10.14806/ej.17.1.200

4.	Callahan BJ, McMurdie PJ, Rosen MJ, Han AW, Johnson AJA, Holmes SP (2016) DADA2: High-resolution sample inference from Illumina amplicon data. Nat Methods 13: 581-583. doi: 10.1038/nmeth.3869

5.	Yilmaz P, Parfrey LW, Yarza P, Gerken J, Pruesse E, Quast C, Schweer T, Peplies J, Ludwig W, Glöckner FO (2014) The SILVA and “all-species living tree project (LTP)” taxonomic frameworks. Nucleic Acids Res 42: D643-D648.

6.	McMurdie PJ, Holmes S (2013) phyloseq: An R package for reproducible interactive analysis and graphics of microbiome census data. PLoS One 8: e61217. doi: 10.1371/journal.pone.0061217

7.	Schloss PD (2021) Amplicon sequence variants artificially split bacterial genomes into separate clusters. mSphere 6: 10.1128/msphere.00191-00121. doi: doi:10.1128/msphere.00191-21

8.	McCauley M, Goulet TL, Jackson CR, Loesgen S (2023) Systematic review of cnidarian microbiomes reveals insights into the structure, specificity, and fidelity of marine associations. Nat Commun 14: 4899. doi: 10.1038/s41467-023-39876-6

9.	Wright ES (2016) Using DECIPHER v2. 0 to analyze big biological sequence data in R. R J 8.

10.	McLaren M (2024) speedyseq: Faster implementations of common phyloseq functions.
